# Fixation matters: duration in fixative prior to immunofluorescent analysis directly impacts macrophage visualisation in epithelial tissues

**DOI:** 10.64898/2026.02.13.705745

**Authors:** Lizi M. Hegarty, Erin Watson, Calum C. Bain, Elaine Emmerson

## Abstract

Macrophages are now recognised as key players in a range of tissues and biological processes, responding to injury and infection, and facilitating development and regeneration. As the importance of macrophage crosstalk within these processes has been revealed, so too has the significance of studying the spatial positioning of macrophages within the tissue of interest. As such, immunofluorescent microscopy-based analysis is becoming an increasingly attractive technique for immunology research. While tissue fixation preserves the tissue architecture and immobilises target antigens, prolonged fixation can negatively impact protein recognition. We report that prolonged exposure to a paraformaldehyde-based fixative profoundly impacts detection of cell surface markers that define macrophage subsets in the mouse submandibular gland, in contrast to epithelial cell markers which appear more robust. We find that this that this is not exclusive to the salivary gland, and similar effects are seen in the pancreas and kidney. Importantly, a short duration of fixation allowed the detection of macrophage subsets in both mouse and human tissue without compromising the detection of other markers. Adoption of a short fixation approach enables accurate detection of a wide range of cell types in tissues, and facilitates exploration of spatial positioning and cell proximity by immunofluorescent microscopy analysis.

## Introduction

Macrophages are innate immune cells performing a spectrum of functions. These include immune surveillance and defence, encompassing specific processes such as phagocytosis, antigen presentation and cytokine secretion (*1*). More recently these functions have been expanded to include a vast range of roles beyond immunity, and macrophages are now also recognised as important regulators of homeostasis and repair, where they contribute by secreting growth factors and mediating metabolism(*2*).

Immunology research has traditionally relied on flow cytometry (*3*) to ascertain the heterogeneity of immune cells within tissues in health, following infection and injury, or during inflammatory resolution or tissue regeneration. However, it is becoming increasingly evident that cellular positioning in the tissue is important (*4-6*) and this spatial data is lost when the tissue is dissociated prior to analysis, such as with flow cytometry. Furthermore, many cells which provide a niche for immune cells in the tissue, such as neurons, do not survive traditional tissue digest and/or are poorly captured by flow cytometry analysis. Thus, many researchers now combine flow cytometry and immunofluorescent microscopy-based analysis for a thorough characterisation of the immune environment.

Immunofluorescent staining has become a fundamental method to detect and visualise a multitude of different cell types *in situ*. The technique involves incubation of the tissue with a primary antibody which recognises and binds to a specific antigen or epitope, followed by detection and amplification by a fluorescently-conjugated secondary antibody, which recognises the host species of the primary antibody. Alternatively, directly-conjugated antibodies provide a one-step staining method. Tissue fixation is an essential step in tissue histology and immunofluorescent staining alike, and acts to not only prevent the tissue from degrading but also to immobilise target antigens and preserve the tissue architecture (*7*). However, excessive fixation can negatively impact protein recognition (*8*), ultimately impacting data interpretation.

We have previously shown that the murine submandibular gland (SMG) harbours at least two distinct macrophage subsets: a CD11c^+^CD163^-^ population and a CD11c^-^CD163^+^ population (*9*). Crucially, we find that these subsets inhabit discrete anatomical locations in the SMG: CD11c^+^CD163^-^ cells associating intimately with epithelial cells, while CD11c^-^CD163^+^ cells appear to associate more with blood vessels and nerves (*9*). The positioning of CD11c^-^CD163^+^ cells in the SMG is in agreement with so-called “TLF” (TIM4 and/or LYVE-1 and/or FRβ–expressing) macrophages, which have been reported across multiple different tissues (*10-12*).

Here, we report that the duration of tissue fixation can profoundly impact the extent of detection of cell surface markers that define macrophage subsets in the mouse SMG. This contrasts with corresponding surface markers on other cell lineages, such as epithelial cell markers which appear more robust and unaffected by the same fixation process. Furthermore, by a comparative analysis of fixation protocols across a number of tissues, we show that this is not exclusive to the SMG, and similar effects are seen in a number of other organs. Thus, it is crucial to ensure that fixation protocols are carefully applied to ensure they are appropriate for the tissue and cell type of interest.

## Materials and Methods

### Animal experiments

All animal experiments adhere to the NC3Rs ARRIVE guidelines and the University of Edinburgh guidelines on the care and use of laboratory animals. All tissues were collected following an approved UK Home Office euthanasia method. We have previously shown that sexual dimorphism exists in mouse SMG structure and immune cells (*9*); thus, for simplicity only male mice were used in this study. Mice were aged between 8 and 12 weeks.

### Human tissue

Human submandibular salivary gland was collected from patients undergoing tissue resection, and from tissue which would otherwise be discarded, under the NHS Lothian BioResource (approval number SR857) and with patient consent.

### Tissue processing

SMGs, kidneys, pancreas and dorsal skin were fixed for either 1, 6 or 24 hours in Antigen Fix (Diapath), a paraformaldehyde-based fixative, at 4 °C, followed by 3 x washes with phosphate buffered saline (PBS; Merck). After fixation, tissue was incubated in 33% sucrose (Sigma Aldrich) overnight at 4 °C before embedding in OCT (Leica). 20 μm sections were cut using a cryostat (Leica) and stored at -20 °C.

### Immunofluorescent analysis

Slides were allowed to come to room temperature. Tissue sections were permeabilised with ice cold acetone/methanol (1:1) for 1 min. Sections were then air dried for 2 minutes, followed by washing in Wash Buffer (PBS + 0.2% BSA) for 3 minutes. Tissue was blocked for 20 minutes at room temperature with Blocking Buffer (1:500 FC Block [Biolegend # 101320] in PBS + 1% BSA). Sections were incubated with primary antibodies overnight at room temperature. Antibodies are listed in **Table 1**. Antibodies were detected using donkey AF488-, Cy3- or AF647-conjugated secondary Fab fragment antibodies (Jackson Laboratories) and nuclei stained using Hoechst 33342 (1:1000, Sigma Aldrich), and mounted using Prolong Gold anti-fade mounting media. Images of 3 random fields of view per sample were acquired on a Leica SP8 confocal microscope. Fluorescent images were collated with NIH ImageJ software and quantified using QuPath software (macrophage cells counts) or Image J (epithelial/endothelial intensity).

**Table 1.** Primary antibodies used for immunofluorescent staining.

| Antibody | Clone | Species | Supplier | Cat # | Dilution | RRID |
| --- | --- | --- | --- | --- | --- | --- |
| AQP5 | EPR3747 | Rabbit | Abcam | ab92320 | 1:200 | AB_2049171 |
| CD11c | N418 | Armenian Hamster | Biolegend | 117302 | 1:100 | AB_313770 |
| CD163 | TNKUPJ | Rat | Thermo Fisher | 14-1631-82 | 1:500 | AB_2716934 |
| CD31 |  | Goat | R&D Systems | AF3628 | 1:200 | AB_2161028 |
| ECAD | ECCD-2 | Rat | Invitrogen | 13-1900 | 1:400 | AB_2533005 |
| IBA1 |  | Rabbit | Antibodies Online | ABIN2857032 | 1:200 |  |

## Results

We have recently demonstrated that macrophages are the predominant immune cell type in the murine submandibular gland (SMG) and that they can be separated into at least two distinct populations: a CD11c^+^CD163^-^ population and a CD11c^-^CD163^+^ population (*9*). However, our early analysis of tissue fixed for 6 hours found that the latter population was minor in comparison to their CD11c^+^CD163^-^ counterparts when assessed by immunofluorescent staining. This was corroborated when we analysed these populations by methods that require tissue dissociation, such as flow cytometry and single cell RNA sequencing (scRNAseq) (*9*). However, this is at odds with findings in other mucosal tissues such as the intestine (*13, 14*) and skin (*15*), where these different macrophage subsets are found in relatively similar abundance. Importantly, there is evidence that CD163^+^ macrophages inhabit different anatomical niches within tissues when compared to their CD163^-^ counterparts (*10*), and as such, being able to accurately locate these cells *in situ* in tissue is becoming increasingly important.

Here, we tested three different durations of tissue fixation and compared expression of some key macrophage and structural cell markers in the SMG. We analysed adult (8-12 weeks of age) male murine submandibular gland (SMG) which had been fixed in Antigen Fix solution (Diapath), a paraformaldehyde-based fixative agent. We compared SMG which had been fixed for 24 hours (i.e. overnight) – a common fixation protocol in salivary gland research (*16-20*) – with two shorter periods of 1 or 6 hours (as used in *21, 22, 23*). We first stained cryosections for the pan-macrophage marker IBA1, and for the subset markers CD11c and CD163. We found that there were no significant differences in the number of IBA^+^ cells between SMG which had been fixed for 1, 6 or 24 hours (**Figure 1A, B**). Conversely, we found that CD163^+^ macrophages in the SMG were abundant after 1 hour of fixation, but the numbers detectable within the tissue decreased after 6 hours of fixation, and were significantly reduced after 24 hours of fixation (**Figure 1A, B**). Similarly, we found that CD11c^+^ macrophages were reliably visible with 1 hour of fixation, and significantly reduced with 6 hours, and even more strikingly after 24 hours fixation (**Figure 1A, B)**. This indicates that while IBA1 staining is robust and detectable across a range of fixation times, expression of CD163 and CD11c in the SMG is notably negatively affected by prolonged fixation of the tissue.

**Figure 1.**
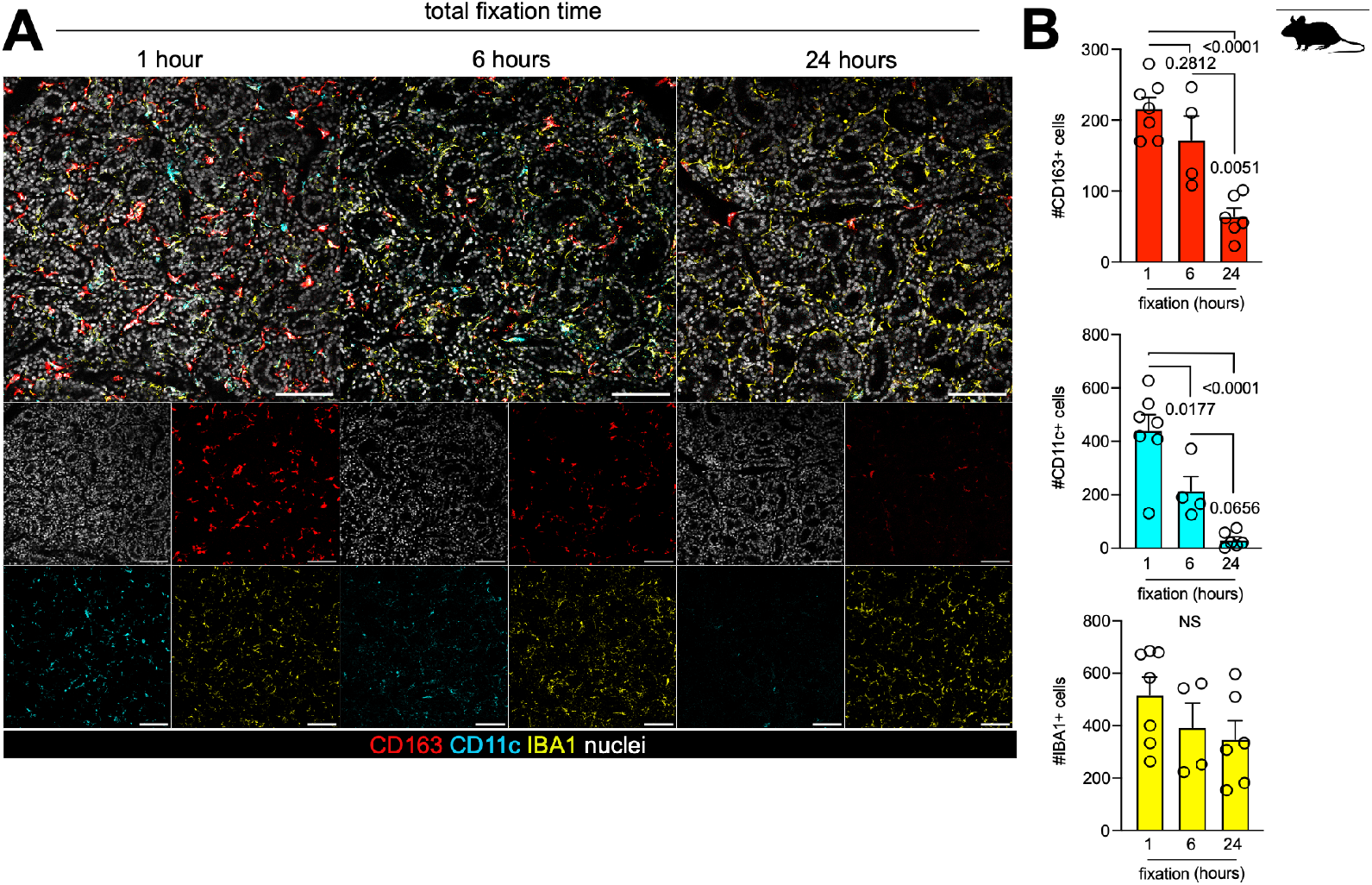
Prolonged tissue fixation impacts macrophage visualisation in the mouse submandibular gland. **A**. Representative expression of CD163, CD11c and IBA1 in SMG tissue from adult male C57BL/6J mice, fixed for 1, 6 or 24 hours. Large images show composites of all channels, insets display individual channels. Scale bar = 100μm. **B**. Graphical enumeration of CD163^+^, CD11c^+^ and IBA1^+^ cells in the mouse SMG at the indicated time points post-fixation. Data was obtained from three fields of view from 4-7 mice per timepoint. Statistical testing was undertaken using one-way ANOVA with Tukey *Q* post-hoc testing. NS = non-significant.

We then extended our analyses to investigate epithelial markers in the SMG. Epithelial integrity and/or expression of epithelial markers is routinely used in salivary gland research to assess tissue function and regeneration/degeneration following injury. Furthermore, macrophages are known to intimately associate with epithelial cells and blood vessels in other organs (*10, 24-26*); and as such, being able to accurately visualise interactions is crucial. We found that the pan-epithelial marker E-cadherin (ECAD) and the acinar cell-specific water channel aquaporin-5 (AQP5) appeared comparable across all fixation conditions (**Figure 2A, C**). This suggests that these epitopes are robust and can be reliably detected regardless of the fixation length. In contrast, while the endothelial cell marker CD31 (PECAM-1) which is often used to mark blood vessels, was apparent after 6 and 24 hours of fixation, the expression was notably stronger after only 1 hour of fixation (**Figure 2B, C**). This implicates over-fixation as a potential cause for concern when assessing the vascularisation of tissue or the nuances of endothelial barrier function.

**Figure 2.**
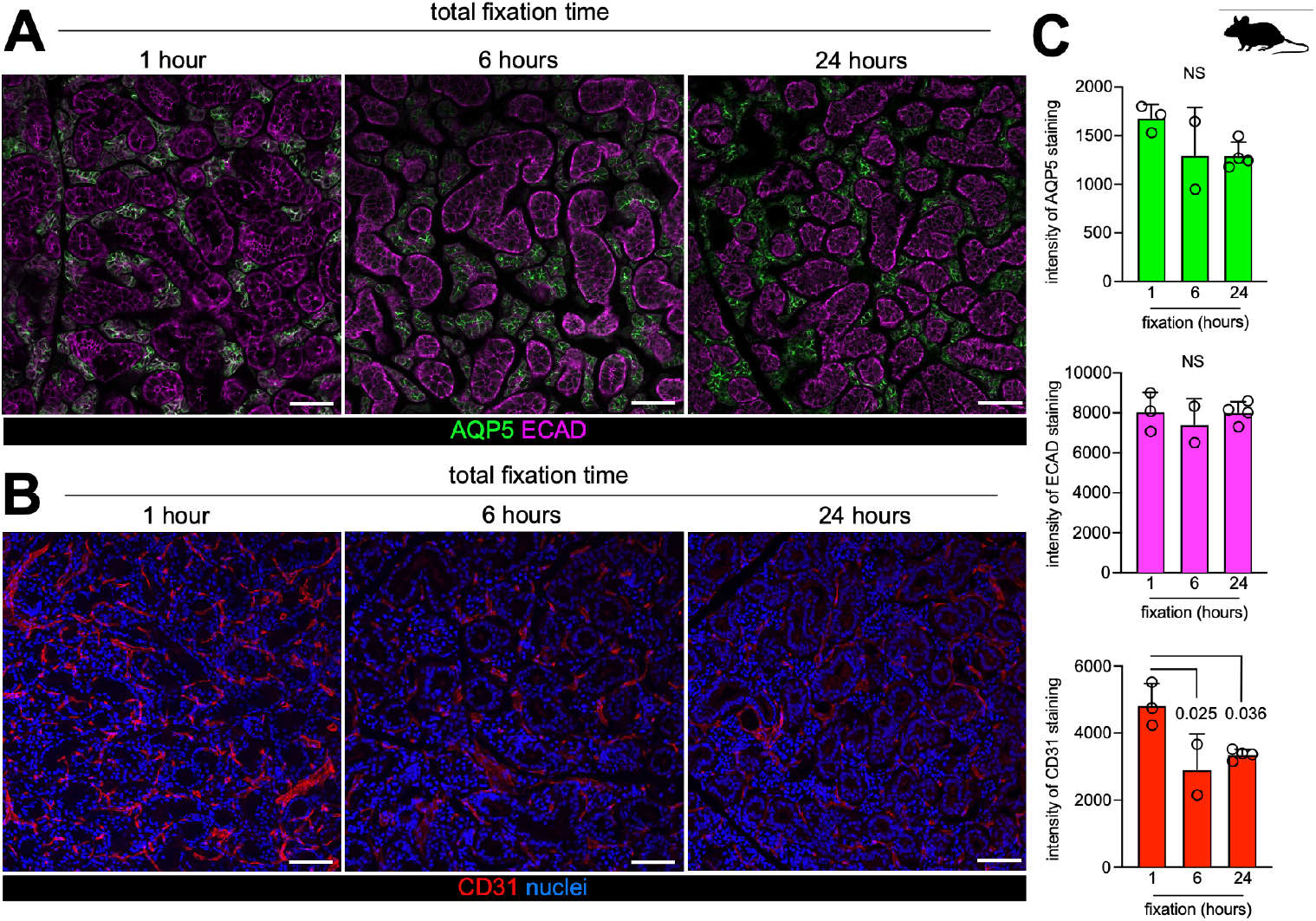
Prolonged fixation impacts submandibular gland endothelial markers but not epithelial markers. **A**. Representative expression of aquaporin-5 (AQP5) and E-cadherin (ECAD) in SMG tissue from adult male C57BL/6J mice, fixed for 1, 6 or 24 hours. Scale bar = 100μm. **B**. Representative expression of CD31 (PECAM-1) in SMG tissue from adult male C57BL/6J mice, fixed for 1, 6 or 24 hours. Scale bar = 100μm. **C**. Graphical enumeration of AQP5, ECAD and CD31 intensity in the mouse SMG at the indicated time points post-fixation. Data was obtained from three fields of view from 2-4 mice per timepoint. Statistical testing was undertaken using one-way ANOVA with Tukey *Q* post-hoc testing. NS = non-significant.

We next set out to find out if the duration of fixation also influenced the visualisation of macrophage subsets in other epithelial tissues. We analysed the mouse pancreas, an organ where roles for macrophages in development and regeneration have recently been described (*27, 28*); the kidney, since this is an organ where an interest in macrophages is flourishing, albeit with limited utility of or success at visualising macrophages by immunofluorescence (*29-31*); and skin, an organ that has long been known to harbour macrophages with roles in repair and regeneration (*32, 33*). Consistent with the SMG, we found that the pancreas, kidney and skin all harbour IBA1^+^, CD11c^+^ and CD163^+^ macrophages (**Figure 3**). Similar to the SMG, while the detection of IBA1^+^ macrophages in the pancreas was unaffected with the increased duration of fixation, expression of CD163 was markedly affected (**Figure 3A**). Intriguingly, CD11c expression was lower in the pancreas than the SMG in all fixation conditions, but did not appear to change with fixation duration (**Figure 3A**). We also analysed mouse kidney, and detected IBA1^+^, CD163^+^ and CD11c^+^ cells, albeit in small number for the latter two (**Figure 3B**). However, the detection of CD163 and CD11c declined after 6 and 24 hours of fixation, when compared to 1 hour (**Figure 3B**). Finally, we analysed mouse dorsal skin fixed for 1, 6 or 24 hours and sectioned in cross-section. Unlike the other epithelial organs that we analysed, we found that CD11c^+^, CD163^+^ and IBA1^+^ cells were all abundant in the dermis after 1 hour of fixation, and their numbers remained consistent after 6- and 24 hours fixation (**Figure 3C**). Collectively, this demonstrates that the duration of fixation negatively impacts visualisation of macrophage subsets in other epithelial organs, in addition to the salivary glands, but that this effect is highly tissue-specific.

**Figure 3.**
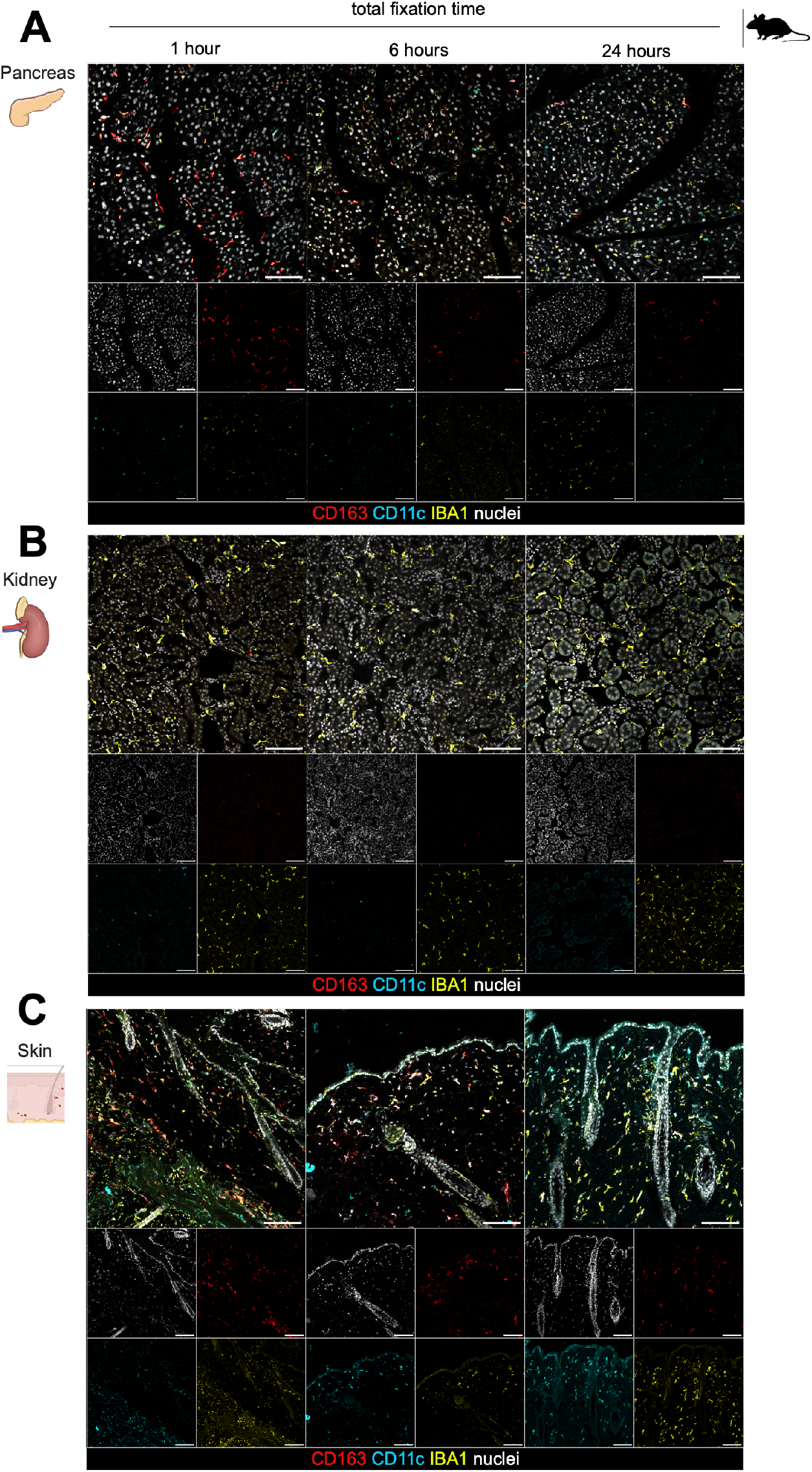
Sensitivity of macrophages to prolonged fixation is tissue-specific. Representative expression of CD163, CD11c and IBA1 in **(A)** pancreas; **(B)** kidney; and **(C)** skin from adult male C57BL/6J mice, fixed for 1, 6 or 24 hours. Large images show composites of all channels, insets display individual channels. Scale bar = 100μm.

In addition, we analysed the pancreas and kidney for the epithelial marker, ECAD, and the endothelial marker, CD31. While expression of ECAD in pancreas was evident across fixation conditions, similar to the SMG, albeit with reduced intensity, fixation of kidney for 6 and 24 hours resulted in clearly disrupted epithelial staining, when compared to 1 hour (**Figure 4A**). Similar to the SMG, CD31 staining was strongest after 1 hour of fixation and expression intensity decreased at 6 and 24 hours (**Figure 4B**). Intriguingly, CD31 expression was not visible in the kidney with any of the fixation conditions analysed (not shown), highlighting the sensitivity of the kidney when analysing by immunofluorescence.

**Figure 4.**
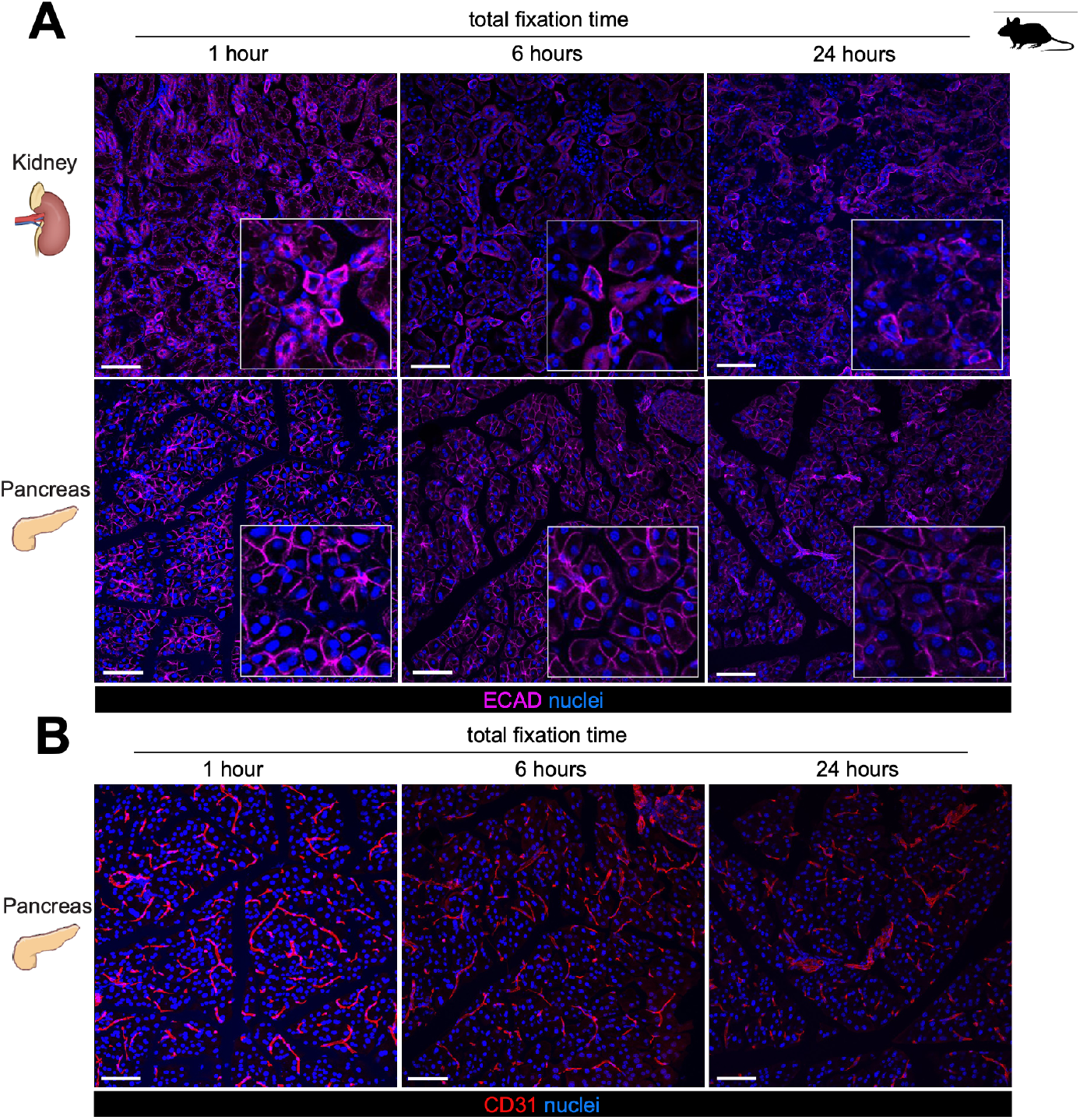
Structural markers are influenced by over-fixation in a tissue-specific manner. **A**. Representative expression of ECAD in kidney and pancreas from adult male C57BL/6J mice, fixed for 1, 6 or 24 hours. Insets display magnified (x4) images. Scale bar = 100μm. **B**. Representative expression of CD31 in pancreas from adult male C57BL/6J mice, fixed for 1, 6 or 24 hours. Scale bar = 100μm.

Finally, we investigated whether we saw a similar sensitivity to fixation time in human salivary gland samples. We analysed human submandibular gland (hSMG) which had been fixed for 1, 6 or 24 hours and stained for IBA1, CD163 and CD11c (**Figure 5**). We found clear CD11c and IBA1 staining in hSMG fixed for 1 hour, while as in mouse SMG, CD11c detection was reduced with 6 and 24 hours. Unlike mouse SMG, IBA1 also appeared to be sensitive to the longer durations of fixation in hSMG, with reduced staining from 6-24 hours fixation (**Figure 5A**). Finally, we found that while epithelial markers were evident under all fixation conditions, AQP5 was more strongly expressed after just 1 hour, when compared to 6 and 24 hours (**Figure 5B**). Importantly, 1 hour of fixation allowed the detection of macrophage subsets in both mouse and human SMG without compromising the detection of other markers, including IBA1, aquaporin-5, E-cadherin and CD31

**Figure 5.**
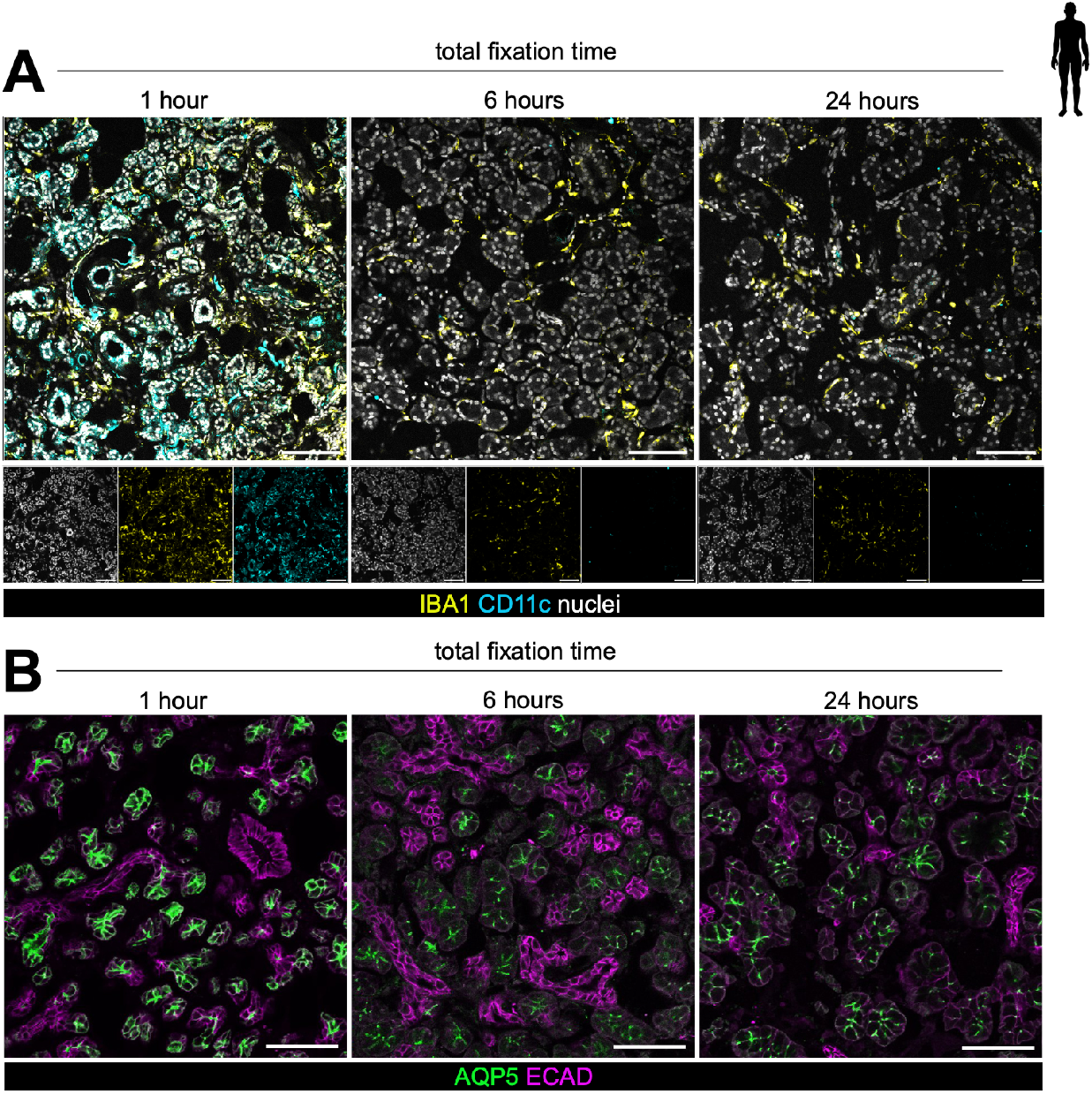
Duration of tissue fixation impacts macrophage visualisation in the human submandibular gland. **A**. Representative expression of CD11c and IBA1 in human submandibular gland (hSMG), fixed for 1, 6 or 24 hours. Scale bar = 100μm. **B**. Representative expression of aquaporin-5 (AQP5) and E-cadherin (ECAD) in human submandibular gland (hSMG), fixed for 1, 6 or 24 hours. Scale bar = 100μm.

Overall, our results demonstrate that the duration of fixation must be carefully considered on a tissue-specific basis whilst taking into account the target proteins of interest, in order to prevent loss of detection and inaccurate conclusions.

## Discussion

Macrophages are now known as key mediators of a multitude of cellular processes beyond their classical phagocytic role, including of processes related to repair and regeneration, amongst others (*2*). Indeed, we have previously shown that macrophages in the submandibular salivary gland (SMG) are crucial in the acute response to irradiation injury (*9*). However, tissue-resident macrophages are not homogenous, and the heterogeneity reported in numerous tissues often relates to their ontogeny, longevity, function and positioning (*34*). Importantly, their expression profile can often inform on their function and anatomical location. For example, border-associated macrophages and microglia in the brain are thought to be distinguishable by their expression, or lack thereof, of CD206, CD163 and CD169 (*35*), markers which are often associated with proximity to blood vessels and scavenging functions. In contrast, IBA1 is considered to be a more generic macrophage marker (*36, 37*). We and others have previously shown that there are at least two transcriptionally distinct subsets of macrophages in the mouse SMG, inhabiting distinct anatomical niches, and which can be separated by their expression of CD11c and CD163 (*9, 16*). These findings are in line with recent studies by the Epelman group, who report that similar macrophage subsets are found across 17 murine tissues (*10*); while other studies have reported macrophage separation on the basis of CD11c and CD163 expression in colon (*13, 14*) and skin (*15*).

In this study we explored whether the duration of tissue fixation impacted the ability to detect these subsets in the SMG by immunofluorescence. We found that while macrophages marked by either CD11c and CD163, were abundant after 1 hour of fixation, this significantly declined when the tissue was fixed for longer periods of 6 or 24 hours, suggesting a sensitivity with prolonged fixation. In contrast, detection of the pan-macrophage marker IBA1, or the epithelial markers aquaporin-5 and E-cadherin, did not appear to change with increased fixation. A number of published studies have aimed to explore aspects of macrophage biology in the mouse submandibular gland using histological techniques. However, many of these studies find a dearth of macrophages in homeostatic conditions when immunofluorescence is used (*16, 38, 39*); a finding which is at odds with other studies which use a combination of methods to assess cellular composition of the tissue (*9, 40, 41*). Thus, in order to ensure that we accurately analyse macrophage abundance and subsets in the future, we have subsequently amended our tissue fixation protocol for immunofluorescent analysis. We find that with a short (1 hour) fixation we preserve both the epitopes and architecture of structural cells (such as epithelial markers) whilst also gaining a more representative view of macrophage abundance and subsets.

Crucially, we found that the sensitivity of these macrophage subsets was not restricted to the salivary gland; and kidney and pancreas also showed a similar propensity to lose the signal of CD11c^+^ and CD163^+^ cells with increased fixation, while skin was visibly more robust. In this study we have not accounted for the fact that cells beyond macrophages, including dendritic cells, monocytes and some T and NK cells, can also express CD11c (*42*). However, by coupling with a pan-macrophage marker, such as IBA1, we are able to make such a distinction using immunofluorescence. While macrophages express different and varying levels of a plethora of markers across different tissues (*43*), our study highlights that fixation has a profound impact on the visualisation of such markers in a range of tissues. Importantly, we find that while some important tissue structures, such as epithelia, are seemingly unchanged by fixation, detection of other structures which provide a niche for tissue macrophages, and vice versa, including the blood vessels (*44-46*), are affected. This is in agreement with reports that paraformaldehyde (PFA) fixation can result in a loss of CD31 signal in brain tissue (*47*), and in line with a recently developed protocol for assessing CD31 in the mouse eye which instead advises fixation for as little as 50 minutes (*48*).

Excessive cross-linking of proteins by prolonged exposure to formalin masks antigenic epitopes, and consequently can prevent antibodies from binding effectively. Overall, our data supports studies which have found that increased fixation impacts the fluorescent signal when analysing oligodendrocyte precursor cells (OPCs), or complement staining in the brain, purportedly due to masking of the antigen (*49-51*). To our knowledge, this is the first report of differences in tissue macrophage abundance with increased fixation. Ultimately, our findings reveal that macrophage visualisation is improved while tissue integrity is not compromised with short fixation; thus, providing an advantage over prolonged fixation. By adopting this tissue processing approach users can accurately quantify a wide range of cell types in tissues, whilst also exploring their spatial positioning and proximity to each other.

## Ethics Approval

*Animal experiments:* All animal experiments adhere to the NC3Rs ARRIVE guidelines and the University of Edinburgh guidelines on the care and use of laboratory animals. All tissues were collected following an approved UK Home Office euthanasia method. EE holds a UK Project Licence (PP0330540).

*Human tissue:* Human submandibular salivary gland was collected under the NHS Lothian BioResource (approval number SR857) with patient consent. Tissue would otherwise be discarded.

## Acknowledgements

The authors would like to thank the Institute for Regeneration and Repair (IRR) Imaging Facility, and the CRM and the Little France (LFR) animal facilities for their technical assistance; and Marc Bajenoff and Johnny Bonnardel for knowledge sharing and critical reading of the manuscript.

## Author Contributions

LMH: Conceptualisation, Data Curation, Formal Analysis, Investigation, Validation, Visualisation, Methodology, Writing – Review & Editing.

EW: Data Curation, Investigation, Methodology, Writing – Review & Editing.

CCB: Conceptualisation, Funding Acquisition, Project Administration, Supervision, Writing – Review & Editing.

EE: Conceptualisation, Data Curation, Formal Analysis, Funding Acquisition, Investigation, Methodology, Project Administration, Resources, Supervision, Validation, Visualisation, Writing – Original Draft, and Writing – Review & Editing.

## References

1. T. Lazarov, S. Juarez-Carreño, N. Cox, F. Geissmann, Physiology and diseases of tissue-resident macrophages. Nature 618, 698–707 (2023).

2. J. Zhao, I. Andreev, H. M. Silva, Resident tissue macrophages: Key coordinators of tissue homeostasis beyond immunity. Sci Immunol 9, eadd1967 (2024).

3. S. P. Perfetto, P. K. Chattopadhyay, M. Roederer, Seventeen-colour flow cytometry: unravelling the immune system. Nature Reviews Immunology 4, 648–655 (2004).

4. M. Guilliams, G. R. Thierry, J. Bonnardel, M. Bajenoff, Establishment and Maintenance of the Macrophage Niche. Immunity 52, 434–451 (2020).

5. D. L. Farber, Tissues, not blood, are where immune cells function. Nature 593, 506–509 (2021).

6. J. M. Schenkel, K. E. Pauken, Localization, tissue biology and T cell state - implications for cancer immunotherapy. Nat Rev Immunol 23, 807–823 (2023).

7. K. Im, S. Mareninov, M. F. P. Diaz, W. H. Yong, An Introduction to Performing Immunofluorescence Staining. Methods Mol Biol 1897, 299–311 (2019).

8. L. Paavilainen, A. Edvinsson, A. Asplund, S. Hober, C. Kampf, F. Pontén, K. Wester, The impact of tissue fixatives on morphology and antibody-based protein profiling in tissues and cells. J Histochem Cytochem 58, 237–246 (2010).

9. J. G. McKendrick, G. R. Jones, S. S. Elder, E. Watson, W. T’Jonck, E. Mercer, M. S. Magalhaes, C. Rocchi, L. M. Hegarty, A. L. Johnson, C. Schneider, B. Becher, C. Pridans, N. Mabbott, Z. Liu, F. Ginhoux, M. Bajenoff, R. Gentek, C. C. Bain, E. Emmerson, CSF1R-dependent macrophages in the salivary gland are essential for epithelial regeneration after radiation-induced injury. Sci Immunol 8, eadd4374 (2023).

10. S. A. Dick, A. Wong, H. Hamidzada, S. Nejat, R. Nechanitzky, S. Vohra, B. Mueller, R. Zaman, Kantores, L. Aronoff, A. Momen, D. Nechanitzky, W. Y. Li, P. Ramachandran, S. Q. Crome, B. Becher, M. I. Cybulsky, F. Billia, S. Keshavjee, S. Mital, C. S. Robbins, T. W. Mak, S. Epelman, Three tissue resident macrophage subsets coexist across organs with conserved origins and life cycles. Sci Immunol 7, eabf7777 (2022).

11. S. Epelman, K. J. Lavine, A. E. Beaudin, D. K. Sojka, J. A. Carrero, B. Calderon, T. Brija, E. L. Gautier, S. Ivanov, A. T. Satpathy, J. D. Schilling, R. Schwendener, I. Sergin, B. Razani, E. C. Forsberg, W. M. Yokoyama, E. R. Unanue, M. Colonna, G. J. Randolph, D. L. Mann, Embryonic and adult-derived resident cardiac macrophages are maintained through distinct mechanisms at steady state and during inflammation. Immunity 40, 91–104 (2014).

12. S. A. Dick, J. A. Macklin, S. Nejat, A. Momen, X. Clemente-Casares, M. G. Althagafi, J. Chen, C. Kantores, S. Hosseinzadeh, L. Aronoff, A. Wong, R. Zaman, I. Barbu, R. Besla, K. J. Lavine, B. Razani, F. Ginhoux, M. Husain, M. I. Cybulsky, C. S. Robbins, S. Epelman, Self-renewing resident cardiac macrophages limit adverse remodeling following myocardial infarction. Nat Immunol 20, 29–39 (2019).

13. L. M. Hegarty, G. R. Jones, A. Biram, C. E. Adams, R. M. Gentek, G. T. Ho, E. Emmerson, C. C. Bain, Tissue resident colonic macrophages persist through acute inflammation and adapt to aid tissue repair. Mucosal Immunol, (2025).

14. I. E. Prise, V. Jayaraman, V. Kästele, R. H. Daw, K. Wemyss, H. Bridgeman, S. Tamburrano, P. Strangward, C. Chew, L. Martens, C. L. Scott, M. Guilliams, A. D. Adamson, J. E. Konkel, T. N. Shaw, J. R. Grainger, CD163 and Tim-4 identify resident intestinal macrophages across sub-tissular regions that are spatially regulated by TGF-β. bioRxiv, 2023.2008.2021.553672 (2023).

15. L. C. Zaba, J. Fuentes-Duculan, R. M. Steinman, J. G. Krueger, M. A. Lowes, Normal human dermis contains distinct populations of CD11c+BDCA-1+ dendritic cells and CD163+FXIIIA+ macrophages. J Clin Invest 117, 2517–2525 (2007).

16. L. Lu, T. Kuroishi, Y. Tanaka, M. Furukawa, T. Nochi, S. Sugawara, Differential expression of CD11c defines two types of tissue-resident macrophages with different origins in steady-state salivary glands. Sci Rep 12, 931 (2022).

17. J. Li, S. Sudiwala, L. Berthoin, S. Mohabbat, E. A. Gaylord, H. Sinada, N. Cruz Pacheco, J. C. Chang, O. Jeon, I. M. A. Lombaert, A. J. May, E. Alsberg, C. S. Bahney, S. M. Knox, Long-term functional regeneration of radiation-damaged salivary glands through delivery of a neurogenic hydrogel. Sci Adv 8, eadc8753 (2022).

18. E. Emmerson, A. J. May, L. Berthoin, N. Cruz-Pacheco, S. Nathan, A. J. Mattingly, J. L. Chang, W. R. Ryan, A. D. Tward, S. M. Knox, Salivary glands regenerate after radiation injury through SOX2-mediated secretory cell replacement. EMBO Mol Med 10, (2018).

19. P. L. Weng, M. H. Aure, T. Maruyama, C. E. Ovitt, Limited Regeneration of Adult Salivary Glands after Severe Injury Involves Cellular Plasticity. Cell Rep 24, 1464–1470.e1463 (2018).

20. K. Zubeidat, Y. Jaber, Y. Saba, O. Barel, R. Naamneh, Y. Netanely, Y. Horev, L. Eli-Berchoer, A. Shhadeh, O. Yosef, E. Arbib, G. Betser-Cohen, C. Nadler, H. Shapiro, E. Elinav, D. J. Aframian, A. Wilensky, A. H. Hovav, Microbiota-dependent and -independent postnatal development of salivary immunity. Cell Rep 42, 111981 (2023).

21. M. Bajénoff, E. Narni-Mancinelli, F. Brau, G. Lauvau, Visualizing early splenic memory CD8+ T cells reactivation against intracellular bacteria in the mouse. PLoS One 5, e11524 (2010).

22. I. Mondor, A. Jorquera, C. Sene, S. Adriouch, R. H. Adams, B. Zhou, S. Wienert, F. Klauschen, M. Bajénoff, Clonal Proliferation and Stochastic Pruning Orchestrate Lymph Node Vasculature Remodeling. Immunity 45, 877–888 (2016).

23. J. Bonnardel, W. T’Jonck, D. Gaublomme, R. Browaeys, C. L. Scott, L. Martens, B. Vanneste, S. De Prijck, S. A. Nedospasov, A. Kremer, E. Van Hamme, P. Borghgraef, W. Toussaint, P. De Bleser, I. Mannaerts, A. Beschin, L. A. van Grunsven, B. N. Lambrecht, T. Taghon, S. Lippens, D. Elewaut, Y. Saeys, M. Guilliams, Stellate Cells, Hepatocytes, and Endothelial Cells Imprint the Kupffer Cell Identity on Monocytes Colonizing the Liver Macrophage Niche. Immunity 51, 638–654.e639 (2019).

24. M. T. Tsai, R. Callaghan, C. Ng, L. Peres-Tintin, D. B. Matthews, N. Stanford, K. Ganesh, M. K. Estes, G. E. Diehl, Intestinal epithelial TLR5 signaling promotes barrier-supportive macrophages. Sci Immunol 11, eadr4057 (2026).

25. A. S. Chikina, F. Nadalin, M. Maurin, M. San-Roman, T. Thomas-Bonafos, X. V. Li, S. Lameiras, S. Baulande, S. Henri, B. Malissen, L. Lacerda Mariano, J. Barbazan, J. M. Blander, I. D. Iliev, D. Matic Vignjevic, A. M. Lennon-Duménil, Macrophages Maintain Epithelium Integrity by Limiting Fungal Product Absorption. Cell 183, 411–428.e416 (2020).

26. L. Smets, M. F. Viola, M. Boesch, J. Raman, L. Van Melkebeke, M. Nobis, E. Flint, N. Pajk, P. Brescia, A. Silvestri, R. Feio-Azevedo, E. Modave, L. De Herdt, A. Sanga, O. T. Pop, O. Govaere, J. Verbeek, A. Denadai-Souza, D. Cassiman, N. Vandamme, B. Boeckx, D. Lambrechts, M. Rescigno, C. Bernsmeier, E. A. V. Jones, J. G. Hengstler, A. Ghallab, C. Scheele, F. Nevens, H. Korf, G. Boeckxstaens, S. W. Van der Merwe, Intestinal blood vessel-associated macrophages and gut-vascular barrier dysfunction in cirrhosis. Gut, (2025).

27. A. Migliorini, S. Ge, M. H. Atkins, A. Oakie, R. Sambathkumar, G. Kent, H. Huang, A. Sing, C. Chua, A. J. Gehring, G. M. Keller, F. Notta, M. C. Nostro, Embryonic macrophages support endocrine commitment during human pancreatic differentiation. Cell Stem Cell 31, 1591–1611.e1598 (2024).

28. A. Grosjean, A. Jalon, C. Leveau, M. Diedisheim, D. A. Bejarano, J. Cuenco, K. Mulder, Z. Liu, A. Le Guernic, M. L. Island, J. Walkiers, G. Ates, C. Materne, A. Goncalves, I. Nemazanyy, L. G. Baudrin, S. Baulande, M. Ropert, J. F. Gautier, A. Hamaï, A. Schlitzer, F. Ginhoux, A. Massie, N. Venteclef, E. Dalmas, An islet-resident macrophage antioxidant program preserves β cell physiology. Sci Immunol 10, eadz5181 (2025).

29. A. S. Puranik, I. A. Leaf, M. A. Jensen, A. F. Hedayat, A. Saad, K. W. Kim, A. M. Saadalla, J. R. Woollard, S. Kashyap, S. C. Textor, J. P. Grande, A. Lerman, R. D. Simari, G. J. Randolph, J. S. Duffield, L. O. Lerman, Kidney-resident macrophages promote a proangiogenic environment in the normal and chronically ischemic mouse kidney. Sci Rep 8, 13948 (2018).

30. C. Chew, O. J. Brand, T. Yamamura, C. Lawless, M. Morais, L. Zeef, I. H. Lin, G. Howell, S. Lui, F. Lausecker, C. Jagger, T. N. Shaw, S. Krishnan, F. A. McClure, H. Bridgeman, K. Wemyss, J. E. Konkel, T. Hussell, R. Lennon, Kidney resident macrophages have distinct subsets and multifunctional roles. Matrix Biol 127, 23–37 (2024).

31. M. D. Cheung, E. N. Erman, K. H. Moore, J. M. Lever, Z. Li, J. R. LaFontaine, G. Ghajar-Rahimi, S. Liu, Z. Yang, R. Karim, B. K. Yoder, A. Agarwal, J. F. George, Resident macrophage subpopulations occupy distinct microenvironments in the kidney. JCI Insight 7, (2022).

32. T. A. Wynn, K. M. Vannella, Macrophages in Tissue Repair, Regeneration, and Fibrosis. Immunity 44, 450–462 (2016).

33. T. Lucas, A. Waisman, R. Ranjan, J. Roes, T. Krieg, W. Müller, A. Roers, S. A. Eming, Differential roles of macrophages in diverse phases of skin repair. J Immunol 184, 3964–3977 (2010).

34. S. Gordon, A. Plüddemann, Tissue macrophages: heterogeneity and functions. BMC Biol 15, 53 (2017).

35. R. Sun, H. Jiang, Border-associated macrophages in the central nervous system. J Neuroinflammation 21, 67 (2024).

36. V. V. Guselnikova, V. A. Razenkova, O. V. Kirik, I. A. Nikitina, V. S. Pavlova, S. I. Zharkina, D. E. Korzhevskii, Detection of Tissue Macrophages in Different Organs Using Antibodies to the Microglial Marker Iba-1. Dokl Biochem Biophys 519, 506–511 (2024).

37. Y. Sasaki, K. Ohsawa, H. Kanazawa, S. Kohsaka, Y. Imai, Iba1 is an actin-cross-linking protein in macrophages/microglia. Biochem Biophys Res Commun 286, 292–297 (2001).

38. Y. Zong, Y. Yang, J. Zhao, L. Li, D. Luo, J. Hu, Y. Gao, L. Wei, N. Li, L. Jiang, Characterisation of macrophage infiltration and polarisation based on integrated transcriptomic and histological analyses in Primary Sjögren’s syndrome. Front Immunol 14, 1292146 (2023).

39. Q. Zhao, S. Pan, L. Zhang, Y. Zhang, A. Shahsavari, P. Lotey, C. L. Baetge, M. A. Deveau, C. A. Gregory, G. M. Kapler, F. Liu, A Salivary Gland Resident Macrophage Subset Regulating Radiation Responses. J Dent Res 102, 536–545 (2023).

40. F. Westermann, S. Tuzlak, V. Kreiner, A. Ignacio, D. Bejarano, M. Bijnen, V. Cecconi, H. van Hove, H. Wang, M. Andreadou, G. Litscher, C. Sparano, R. Fróis-Martins, A. Gallerand, E. Roussel, L. Oberbichler, R. Lindemann, D. DeFeo, Z. Liu, A. Kipar, S. LeibundGut-Landmann, K. McCoy, I. Nixon, C. C. Bain, C. Schneider, S. Ivanov, S. Tugues, M. Greter, F. Ginhoux, A. Schlitzer, E. Emmerson, B. Becher, Adenophages are an atypical macrophage population in exocrine glands sustained by ILC2-derived GM-CSF. Nat Immunol 27, 26–34 (2026).

41. B. Stolp, F. Thelen, X. Ficht, L. M. Altenburger, N. Ruef, V. Inavalli, P. Germann, N. Page, F. Moalli, A. Raimondi, K. A. Keyser, S. M. Seyed Jafari, F. Barone, M. S. Dettmer, D. Merkler, M. Iannacone, J. Sharpe, C. Schlapbach, O. T. Fackler, U. V. Nägerl, J. V. Stein, Salivary gland macrophages and tissue-resident CD8(+) T cells cooperate for homeostatic organ surveillance. Sci Immunol 5, (2020).

42. L. Hou, S. Koutsogiannaki, K. Yuki, Multifaceted, unique role of CD11c in leukocyte biology. Front Immunol 16, 1556992 (2025).

43. B. Chen, R. Li, A. Kubota, L. Alex, N. G. Frangogiannis, Identification of macrophages in normal and injured mouse tissues using reporter lines and antibodies. Sci Rep 12, 4542 (2022).

44. K. R. Mesa, K. A. O’Connor, C. Ng, S. P. Salvatore, A. Dolynuk, M. R. Lomeli, D. R. Littman, Niche-specific dermal macrophage loss promotes skin capillary ageing. Nature 648, 173–181 (2025).

45. I. Mondor, M. Baratin, M. Lagueyrie, L. Saro, S. Henri, R. Gentek, D. Suerinck, W. Kastenmuller, J. X. Jiang, M. Bajénoff, Lymphatic Endothelial Cells Are Essential Components of the Subcapsular Sinus Macrophage Niche. Immunity 50, 1453–1466.e1454 (2019).

46. D. C. Saunders, K. I. Aamodt, T. M. Richardson, A. J. Hopkirk, R. Aramandla, G. Poffenberger, R. Jenkins, D. K. Flaherty, N. Prasad, S. E. Levy, A. C. Powers, M. Brissova, Coordinated interactions between endothelial cells and macrophages in the islet microenvironment promote β cell regeneration. NPJ Regen Med 6, 22 (2021).

47. A. C. T. Teng, D. Mehangrey, A. Vandenbelt, K. Vearncombe, J. D. Callahan, P. Mistry, W. Li, C. J. Reitz, O. Hamed, M. Roche, U. Kuzmanov, J. E. Fish, S. Epelman, A. O. Gramolini, Glyoxal is a superior fixative to formaldehyde in promoting antigenicity and structural integrity in murine cardiac tissues. J Mol Cell Cardiol Plus 12, 100454 (2025).

48. W. Wang, P. Peng, W. Hu, X. Liu, L. Yang, R. Ju, F. Zhang, Protocol for immunofluorescent staining, high-resolution imaging, and spatial projection of Schlemm’s canal in mouse whole-mount corneal limbus tissue. STAR Protoc 6, 103906 (2025).

49. F. Pfeiffer, A. Sherafat, A. Nishiyama, The Impact of Fixation on the Detection of Oligodendrocyte Precursor Cell Morphology and Vascular Associations. Cells 10, (2021).

50. T. Mori, T. Wakabayashi, Y. Takamori, K. Kitaya, H. Yamada, Phenotype analysis and quantification of proliferating cells in the cortical gray matter of the adult rat. Acta Histochem Cytochem 42, 1–8 (2009).

51. N. Scott-Hewitt, M. Mahoney, Y. Huang, N. Korte, T. Yvanka de Soysa, D. K. Wilton, E. Knorr, K. Mastro, A. Chang, A. Zhang, D. Melville, M. Schenone, C. Hartigan, B. Stevens, Microglial-derived C1q integrates into neuronal ribonucleoprotein complexes and impacts protein homeostasis in the aging brain. Cell 187, 4193–4212.e4124 (2024).

